# Transcriptome comparison between the cultured and in vivo Chick Primordial Germ Cells by SMART-seq-based single cell RNA sequencing

**DOI:** 10.1101/2025.07.02.662725

**Authors:** Yoshiki Hayashi, Atsushi Doi, Hiroko Iikawa, Hirokazu Kimijima, Yutaka Suzuki, Akinori Kanai, Hideki Hirakawa, Daisuke Saito

**Affiliations:** Graduate School of Science, Kyushu University, 744, Motooka, Nishi-ku, Fukuoka, 819-0395, Japan; Medical Institute of Bioregulation, Kyushu University, 3-1-1, Maidashi, Higashi-ku, Fukuoka, 812-8582, Japan; Graduate School of Systems Life Sciences, Kyushu University, 744, Motooka, Nishi-ku, Fukuoka, 819-0395, Japan; Graduate School of Biostudies, Kyoto University, Yoshida-Konoecho, Sakyo-ku, Kyoto, 606-8501, Japan; Life Science Data Research Center, Graduate School of Frontier Science, The University of Tokyo, 5-1-5, Kashiwanoha, Kashiwa, Chiba, 277-8561, Japan; Graduate School of Bioresource and Bioenvironmental Science, Kyusyu University, 744, Motooka, Nishi-ku, Fukuoka, 819-0395, Japan

**Author notes:** Correspondence: Yoshiki Hayashi, Daisuke Saito, Graduate School of Science, Kyushu University, 744, Motooka, Nishi-ku, Fukuoka, 819-0395, Japan.

**Keywords:** Primordial Germ Cell, Transcriptome, Chick

## Abstract

Primordial germ cells (PGC), the precursors of the germline, have unique cellular characteristics to undergo long-distance migration to the embryonic gonads and have the potential to differentiate into somatic cells. Among the animal models studying PGC development, the chicken PGCs are an ideal model, since it is a rare model in which long-term PGC cultivation is applicable. Although the cultural applicability of chicken PGC makes it attractive for revealing the PGC character and its developmental processes, some differences from endogenous PGCs are known, such as the remarkable up-regulation of cell proliferation and a lesser ability to reach the gonads. Understanding these differences at the molecular level is crucial. To this end, we first performed SMART-seq-based single-cell RNA sequencing to compare transcriptomes between endogenous PGCs and cultivated PGCs. Our results revealed that PGC cultivation causes a shift from a MYC-dependent to a MYCN-dependent gene regulatory network in PGCs, suggesting that this reprogramming contributes to the acquisition of proliferation ability and stem cell characteristics in cultivated PGCs. Additionally, our results suggest that the MYCN-dependent gene regulatory network increases the risk of somatic differentiation, particularly in neural fate, in cultivated PGCs. In addition, our transcriptome analysis identified new cell populations that show molecular character as intermediate cell states between germline and pluripotent cells from the early embryonic stage. Thus, our study provides fundamental molecular information to understand both the effects of PGC cultivation and the developmental process of chicken PGCs.

## Introduction

PGCs are the precursors of the germline and contribute to the life continuity of the species. In addition to their importance as the precursors of the cells in charge of transferring the inheritable materials to the next generation, PGCs have unique cellular and developmental characteristics rarely observed in other cell types in the somatic tissues. First, PGCs of some animal species, including insects and vertebrates, have been proven to have the potential to differentiate into somatic cells under some conditions (Hayashi *et al*., 2004, Gross-Thebing *et al*., 2017, Youngren *et al*., 2005). Thus, PGCs are potentiating to have stem cell-like characteristics. The second discriminating characteristic of PGCs is the ability to migrate long distances within the embryo (Richardson & Lehmann, 2010). PGCs in most animal species studied are formed at early embryogenesis outside of the gonads, and migrate through several different circumstances to reach the embryonic gonads, where they eventually differentiate into gametes. During migration, PGCs need to maintain their germline fate with their fragile character as undifferentiated cells within the complicated developmental environment, and at the same time, they need to precisely accept migration cues from the outside to reach the gonads.

Thus, PGCs are an ideal cellular model to investigate the mechanisms balancing the maintenance of cellular character and response to environmental signals. Avian PGCs, especially, provide a valuable model to address those matters for several reasons. First, Avian PGCs undergo migration via a unique route, the blood vessel (Nakamura *et al*., 2007, Saito *et al*., 2022, Ichikawa & Horiuchi, 2023). Chicken PGCs, as an example, are localized in the central zone of the area pellucida, on the ventral surface of the epiblast (Embryonic day 0; E0). Those PGCs migrate anteriorly to the germinal crescent region where they enter the embryonic circulation associated with the formation of the blood vascular network at E1.25(Murai *et al*., 2021). After circulation, PGCs arrested in the capillaries near the germinal epithelium undergo extravasation to be outside the vasculature at around E2.5(Saito *et al*., 2022, Ando & Fujimoto, 1983). PGCs then migrate along the dorsal mesentery to reach the embryonic gonads (E4.5). Thus, Avian PGCs undertake complicated cancer-like behavior to reach the gonads; they must proceed through intra- and extravasation for precise homing to the gonads.

The second merit of using chicken PGCs is the technical aspects; Chicken PGCs are the first and few PGCs that can be applied to long-term culture after primary isolation from the embryos (Whyte *et al*., 2015, van de Lavoir *et al*., 2006). Those cultured PGCs are easily propagated and are also able to produce the next generation when they are injected back into the embryos. In addition, those cultured PGCs are applicable for transfection of plasmid, thus suitable for analyzing the gene function involved in migration processes (Saito *et al*., 2022, Suzuki *et al*., 2023). Therefore, chicken PGCs provide an ideal model for understanding PGC characteristics and the molecular mechanisms regulating in vivo hematogenous migration processes in general. Although the cultivated PGC (cPGCs) is a useful tool, it has been known that the cPGCs have several different characteristics not observed in endogenous PGCs (ePGCs). First, cPGCs have a stronger proliferation ability than ePGCs; cPGCs easily propagate to x10^6^ order number within a few weeks from a few hundred of their sourced endogenous PGCs (Whyte *et al*., 2015, Song *et al*., 2014, Nakamura *et al*., 2007). Second, cPGCs have less ability to be integrated into embryonic gonads than ePGCs; even when 10 times more cPGCs are injected back into the embryo, the contribution of the ePGCs to generate descendants is limited, suggesting that cPGCs are less competitive to reach the gonads than ePGCs (Song *et al*., 2014). This disability of cPGCs to make the next generation leads to innovative overcoming experimental techniques in which ePGCs are experimentally ablated from the host embryos, so that cPGCs have no competition with ePGCs for reaching the gonads (Trefil *et al*., 2017, Nakamura *et al*., 2010, Nakamura *et al*., 2012, Woodcock *et al*., 2019, Chen *et al*., 2023). Thus, even though cPGCs are a useful model, it is fundamentally important to understand the difference between cPGCs and ePGCs for future use of cPGCs in cellular and/or developmental PGC models.

To understand those characters, basic information about gene expression is essential. To date, several studies provide genome-wide gene expression information about Chicken PGCs. For example, gene expression profiles of ePGCs are developmentally analyzed with bulk-RNA sequencing (RNA-seq) and single-cell RNA-seq (scRNA-seq)(Rengaraj *et al*., 2022a, Gong *et al*., 2024, Rengaraj *et al*., 2022b, Ichikawa *et al*., 2022). Also, gene expression profiles of cPGCs are analyzed with RNA-seq and scRNA-seq (Gong *et al*., 2024, Zou *et al*., 2025). Although those studies provide important basic information to understand the differences between the ePGCs and cPGCs, there are several matters to consider when comparing the data. For example, those studies were performed independently with different PGC sources, genetic backgrounds, and sequencing platforms. In addition, about the scRNA-seq, all the above studies utilized 10x Chromium-based library construction, which is known to have cost-effective performance but provides relatively less effective detection of gene expression. Thus, obtaining deep and fine gene expression profiles from comparative sources is important to understand the differences between ePGCs and cPGCs.

In this study, we performed SMART-seq-based scRNA-seq to compare the gene expression profile between ePGCs and cPGCs from the same laboratory condition. Although less cost-effective and with a limitation on the sample number, our SMART-seq-based scRNA-seq successfully showed high detectability for gene expression from each single PGC. Our gene regulatory network (GRN) analysis revealed that the MYC-dependent transcriptional network was the most affected target of PGC cultivation, and remodeling from MYC- to MYCN-dependent network is achieved during the establishment of cultivated PGCs. Also, our results suggest that the MYCN-dependent gene regulatory network increases the risk of somatic differentiation, particularly in neural fate, in cultivated PGCs. Finally, we provide molecular evidence suggesting that our previously developed methods of collecting PGCs based on Tomato-lectin (Iikawa *et al*., 2024) are useful tools for capturing PGCs from the earliest embryonic stage. Thus, our study provides the fundamental molecular knowledge to understand the effects of PGC cultivation and to help us further understand PGC specification processes in avian species.

## 2. Results & Discussion

### 2.1 Single-cell gene expression profiling of ePGCs and cPGCs

To evaluate the effects of PGC cultivation, we collected 96 PGCs from embryos at E2.5, which is the source of PGC cultivation (ePGC_E2.5), and 48 cultivated PGCs (cPGCs) from each of initiating period of PGC cultivation (1 week after endogenous PGCs are in culture medium; cPGC_1w) or period in which the cultivated PGCs are well-established (3 weeks after PGCs are in culture medium; cPGC_3w), by using our previously developed methods for efficient collection of PGCs: Lycopersicon Esculentum (Tomato) lectin (LEL) based labeling of PGCs combined with Fluorescence-Activated Cell Sorting (FACS)(Iikawa *et al*., 2024). Our LEL-labeling methods have been shown to effectively collect PGCs from E2.5 embryos compared to the method that depends on the PGC-labeling with SSEA1, which is a commonly used labeling method of PGCs (Jung *et al*., 2005). Previously, we showed that LEL-labeled PGCs consistently expressed the evolutionarily conserved PGC markers DEAD-box helicase 4 (DDX4) and Deleted in azoospermia-like (DAZL), but approximately half of LEL-labeled PGCs are not labeled with SSEA1, indicating that the LEL-labeling method is more efficient in collecting PGCs than the SSEA1 method. Also, our previous work showed that LEL-methods label morphologically indistinguishable PGC-like cells from E0 stage embryos, which are absent from SSEA1. Although those results indicate the possibility that our LEL-method could collect the PGCs from more prior embryonic stages than the SSEA1-methods, the lack of molecular information of those PGC-like cells prevents confirming the efficiency of the LEL-method on this point. Thus, we determined to investigate the gene expression of those PGC-like cells additionally to evaluate the possibility that our LEL method could collect PGCs from the earliest embryonic stage. To this end, we additionally collected 99 LEL-positive PGC-like cells (ePGC_E0) and performed SMART-based single-cell RNA sequencing. We sequenced all of the above libraries and evaluated the sample quality based on the number of genes detected. The number of genes detected from most of the samples was more than 12000 genes (Figure 1A), which is obviously a higher number than 10x chromium-based scRNA-seq, which detects a few thousand genes from chick PGCs [such as in (Zou *et al*., 2025)], indicating the usefulness of our SMART-seq-based scRNA-seq. We set our sample cut-off to samples with 100 genes per sample, and as a result, we obtained a total of 162 samples for further analysis. Uniform manifold approximation and projection (UMAP) analysis showed our samples were clustered in four groups, with those roughly merged with their sample source, except ePGC_E0 (Figures 1B, C). Samples from ePGC_E0 were divided into two clusters (cluster 0 and 3 in Figure 1B), one of which was a cluster whose main constituent is ePGC_3w samples (Cluster 0, Figures 1B, C). We confirmed the expression amplitude of DAZL among those samples and found that DAZL expression is the highest in cluster 0, and most of the samples in other clusters have moderate expression of DAZL, except cluster 3, which shows the lowest or absent expression of DAZL (Figure 1D). Those results showed that two populations of PGC-like cells from E0 embryos in different clusters have different characteristics for DAZL expression. In the following analysis, we referred to the names of each cluster as the main constituents of each cluster (Cluster 0 as cluster cPGC_3w_mix, Cluster 1 as ePGC_E2.5, Cluster 2 as cPGC_1w, Cluster 3 as ePGC_E0, Figure 1C).

**Figure 1.**
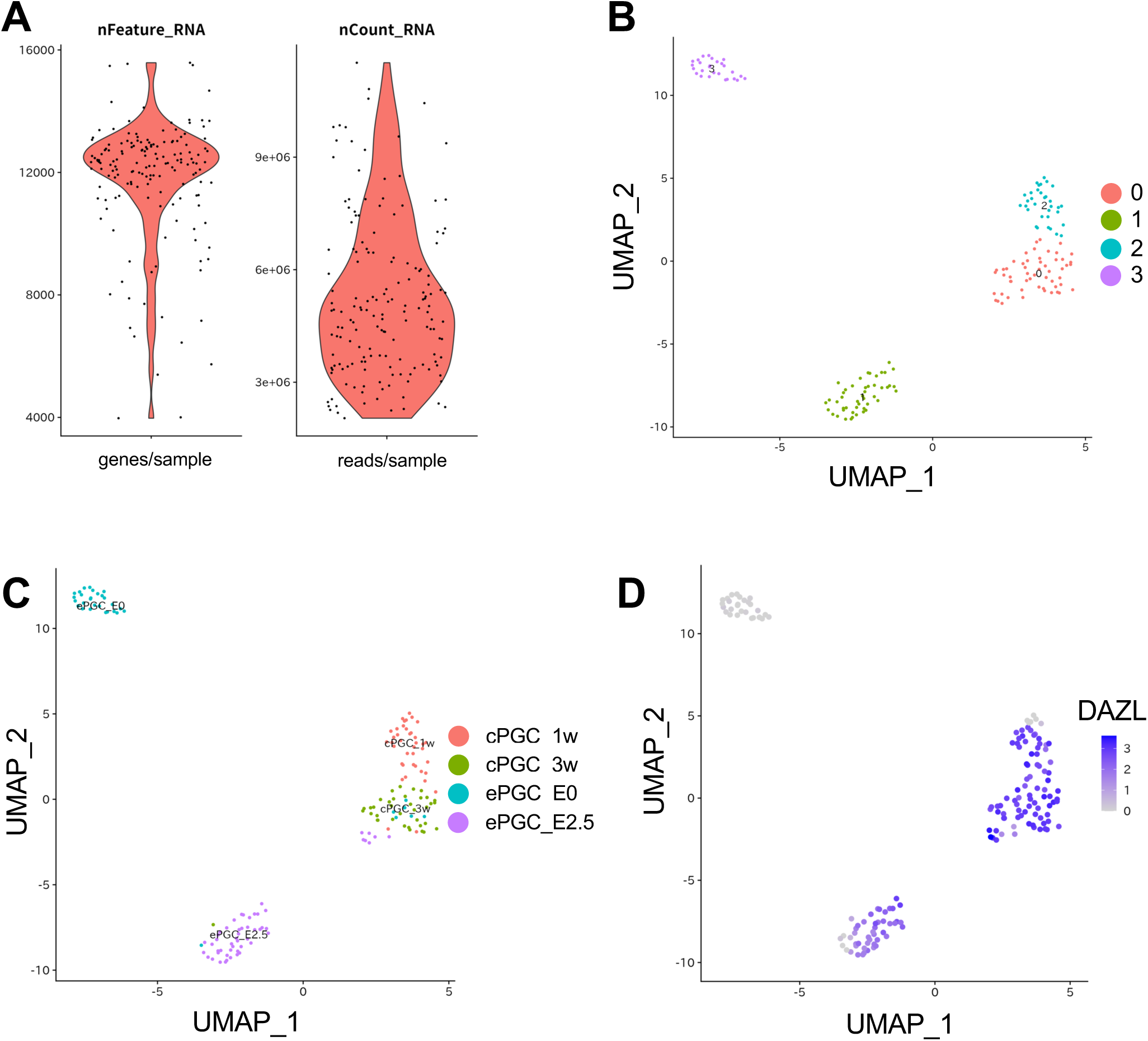
**Single-cell gene expression profiling of ePGCs and cPGCs**. Gene expression profiles of endogenous PGCs and cultivated PGCs were shown. (A) The Distribution of detected gene number (Left) and number of obtained reads (Right) from each cell is shown. (B, C) The Results of UMAP-based clustering analysis are shown. Our total samples were clustered into 4 clusters (B), with constituents of each cluster coming from the same sample category, except cluster 0 (C). (D) The Expression of DAZL, the germline marker, in each cell is shown. Color value is based on normalized log-transformed value.

### 2.2 Molecular feature of each PGC cluster

To investigate the gene expression character for each cluster, we identified marker genes of each cluster; genes whose expressions are significantly up-regulated in one cluster compared to the others. Our clustering analysis of gene expression from each cluster successfully identified marker genes of each cluster (number of marker genes; cPGC_3w_mix: 2366 genes, ePGC_E2.5: 608 genes, cPGC_1w: 476 genes, ePGC_E0: 1412 genes. Top 10 marker genes are shown in Figure 2A, Supp. table 1). To see the overall molecular trend of each cluster, we performed Gene Ontology (GO) enrichment analysis and KEGG pathway analysis using marker genes (Figures 2B-E, Supp. Figure 1). Our GO enrichment analysis showed that ePGC_E2.5, which is the source of cPGCs, has up-regulation of genes with GO (BP: Biological Processes) related to heme and its constituents (Figure 2B) (porphyrin, belonging to the tetrapyrrole group, is a constituent of heme). Also, marker genes of ePGC_E2.5 showed enrichment of GO(BP) related to oxygen metabolism (reactive oxygen species metabolic process and oxygen transport). ePGC_E2.5-marker genes also have mitochondria-related GOs in the GO categories (MF: Molecular Functions, CC: Cellular Components) (Supp. Figures 1A, B). These enriched GOs were related to mitochondrial activity, suggesting that PGCs of embryonic day 2.5 have higher mitochondrial activity than other conditions. GO enrichment analysis for cPGC_1w was less informative, but KEGG pathway analysis for those marker genes showed that mitophagy and ferroptosis pathways were up-regulated in this cell population (Figure 2C), suggesting that removal of mitochondrial and causal release of iron ions occur during the first step of PGC cultivation. It is possible that metabolic reprogramming from mitochondrial oxidative phosphorylation (OXPHOS) to glycolysis, which is frequently observed in proliferating cells (DeBerardinis *et al*., 2008), occurs in the initiation step of PGC cultivation. GO enrichment analysis for cPGC_3w_mix marker genes showed most of the GO (BP)s enriched in this population were related to cell morphology, especially to the formation of cellular projections, such as neuron projection (Figure 2D). Even though this cell population stably expressed germline marker DAZL (Figure 1D), the enriched GOs suggest that cellular differentiation processes were partially enhanced in this cell population. Those results suggest that established PGCs partially possesses the molecular characteristics of other cell lineages, such as neuronal cell lineage. Finally, we performed GO enrichment analysis for the ePGC_E0 cluster (Figure 2E), which is absent for DAZL expression (Figure 1D). GO enrichment analysis for this cell population revealed that GO (BP)s related to translation (e.g., ribosome biogenesis) and post-transcriptional regulation (e.g., ribonucleoprotein complex biogenesis) were enriched in marker genes of the ePGC_E0 cluster. GO enrichment analysis of other GO categories (MF and CC) and KEGG pathway analysis also suggests the enhancement of translational regulation in this cell population. Since the regulation of maternally inherited RNA is critical in cells in early embryonic stages, these results might reflect the cellular status of those developmental situations.

**Figure 2.**
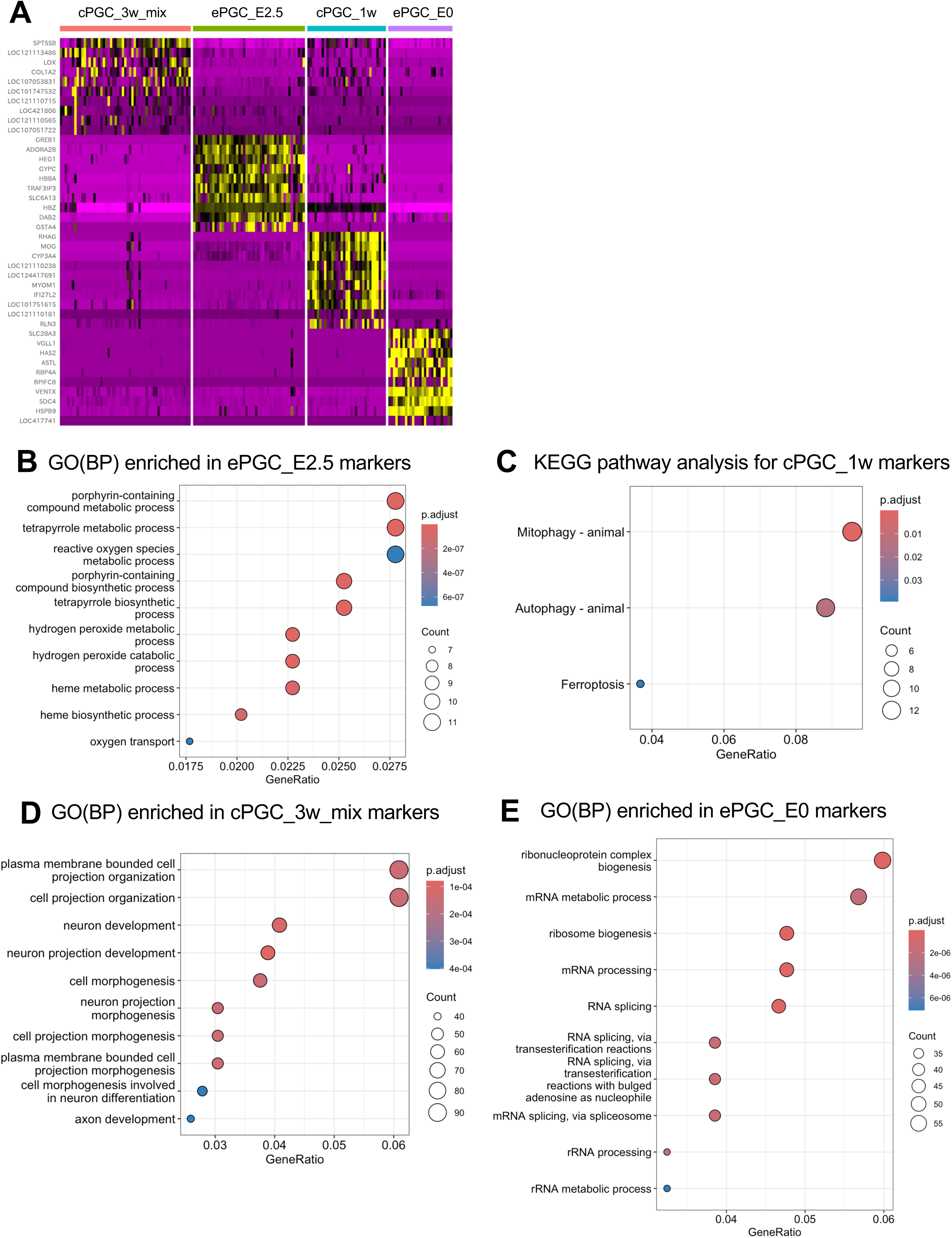
**Molecular feature of PGC clusters**. (A) The heat map of the top 10 marker genes for each cluster is shown. The labels on the top show the original clusters, and the labels on the left indicate the names of marker genes. (B-E) The results of Gene Ontology (GO) enrichment analysis or KEGG pathway analysis for marker genes of each cluster are shown. The analyzed marker gene source and type of analysis are indicated at the top of each figure (e.g., GO(BP) enriched in ePGC_E2.5 markers means the results of GO enrichment analysis for Biological Process (BP) for marker genes of the ePGC_E2.5 cluster).

### 2.3 Gene expression comparison between ePGCs and cPGCs

Next, we determined to compare the gene expression profile of ePGCs and cPGCs in depth to evaluate the effects of PGC cultivation. To this end, we directly compared the gene expression profiles between ePGC_E2.5 and cPGC_1w based on the sample origin, not the clusters. We detected a total of 1148 genes that were differentially expressed in cPGC_1w (747 up-regulated genes and 401 down-regulated genes) compared to ePGC_2.5 (Figure 3A). We found that one of the top down-regulated genes in cPGC_1w was MYC (Figure 3A, approximately 40-fold up-regulation, Supp. Table 2), which is the well-known transcriptional factor (TF) involved in regulation of cell pluripotency and cell proliferation (Whitfield & Soucek, 2025, Bretones *et al*., 2015, Chappell & Dalton, 2013, Kim *et al*., 2010). The gene regulatory network (GRN) analysis between TFs and target genes for those differentially expressed genes (DEGs) showed that MYC is a top hub TF in that GRN (Figure 3C, Supp. Table 3), emphasizing the importance of downregulation of MYC-dependent GRN in initial period of PGC cultivation. About the target genes, we observed up-regulation of CDKN1A/p21, which is cyclin dependent kinase inhibitor to repress the cell cycle (Karimian *et al*., 2016), and is known to be a repressive target of MYC (Bretones *et al*., 2015, Wu *et al*., 2003); Those observations suggest that MYC-dependent GRN works in PGCs. Interestingly, GATA6, which is the endoderm master regulator and key the repressive targets of MYC in pluripotent stem cells are up-regulated in this condition (Figure 3C)(Smith *et al*., 2010), suggesting that PGCs in initial cultivation period slightly loose their stem cell like character. Of note, the SOX2, the transcriptional factor regulating cell pluripotency was also down-regulated in cPGC_1w (Figure 3C). Thus, our transcriptome comparison strongly suggests that the loss of stem cell-like character of ePGCs occurs in the initial step of PGC cultivation. MYC is well known as a repressive target of Smad3(Chen et al., 2002); the downstream signal transducer of Activin A, which is one of the three constituent ligands of PGC cultivation medium (Whyte *et al*., 2015). Although the signaling pathways regulated by both of other two of three constituent ligands, Insulin and FGF2 are known to be involved in up-regulation of MYC(Sears *et al*., 2000, Lee *et al*., 2022), our results suggest that Activin A-dependent TGF-beta signaling has strong effects on the PGC state alternation occurring in the initial step of PGC cultivation. Next, we compared the gene expression profiles between ePGC_E2.5 and cPGC_3w, and between cPGC_1w and cPGC_3w. We detected a total of 2459 genes that were differentially expressed in cPGC_3w (1963 up-regulated genes and 496 down-regulated genes) compared to ePGC_2.5 (Figure 3B, Supp. Table4), and a total of 635 genes (292 up-regulated and 343 down-regualted) compared to cPGC_1w (Supp. Figure 2A, Supp. Table6). As in the case of cPGC_1w, MYC was significantly down-regulated in cPGC_3w (approximately 24-fold upregulation, Supp. Table 4). Our GRN analysis on DEGs based on TFs and their known targets revealed that MYC was the first top hub TFs in the GRN of DEGs (Figure 3D, Supp. Table 5), suggesting that MYC-dependent GRN still affected by cultivation. Interestingly, we found that expression of MYCN, a MYC family gene which has functional and structural similarity to MYC (Chappell & Dalton, 2013), is up-regulated in cPGC_3w (Figure 3D, supp. table 4). Up-regulation of MYCN in cultivated PGCs occurs during the cultivation, since transcriptome comparison between cPGC_1w and cPGC_3w detects MYCN as a differentially expressed gene (Supp. Figure 2A, Supp. Table 6). The GRN analysis on DEGs between cPGC_1w and cPGC_3w showed that MYCN-dependent network formed (Supp. Figure 2B). We found down-regulation of CDKN1A/p21 occurs during the cultivation period between 1 week and 3 weeks (Supp. Figure 2B), suggesting that re-acquisition of proliferation ability could occur during this period, probably in a MYCN-dependent manner (Supp. Figure 2B). MYCN is known to be involved in the regulation of cellular pluripotency, similar to MYC (Chappell & Dalton, 2013). Interestingly, we observed up-regulation of OCT4/ Pou5f3 and NANOG occur in cPGC_3w (Figure 3D, Supp. Table4, Supp. Table6). Conversely, we found GATA6 is down-regulated in cPGC_3w (Figure 2B, Supp. Table6). Those observations strongly suggest that PGCs cultivated for 3weeks re-acquire the proliferation ability and pluripotent character in a MYCN-dependent manner. It has been shown that the forced expression of MYCN in chicken neural crest cells induces CNS-like neural stem cell characteristics in these cells(Kerosuo *et al*., 2018). Indeed, enrichment analysis of DEGs between the cPGC_1w and cPGC_3w using the ARCHS4_Tissues database revealed that DEGs were significantly enriched for those highly expressed in neural cells (Supp. Table 7), suggesting that cultivated PGCs partially possess the neural cell character in gene expression level. This observation is consistent with the GO enrichment analysis for marker genes of cluster cPGC_3w_mix shown above. Therefore, our results suggest that shifting to MYCN-dependent GRN in cultivated PGCs contributes to the maintenance of cell proliferation and stem cell character, but at the same time, it induces neural cell-like character in those. This phenomenon could account for why the cultivated PGCs show less competitiveness to reach the embryonic gonads compared to endogenous PGCs.

**Figure 3.**
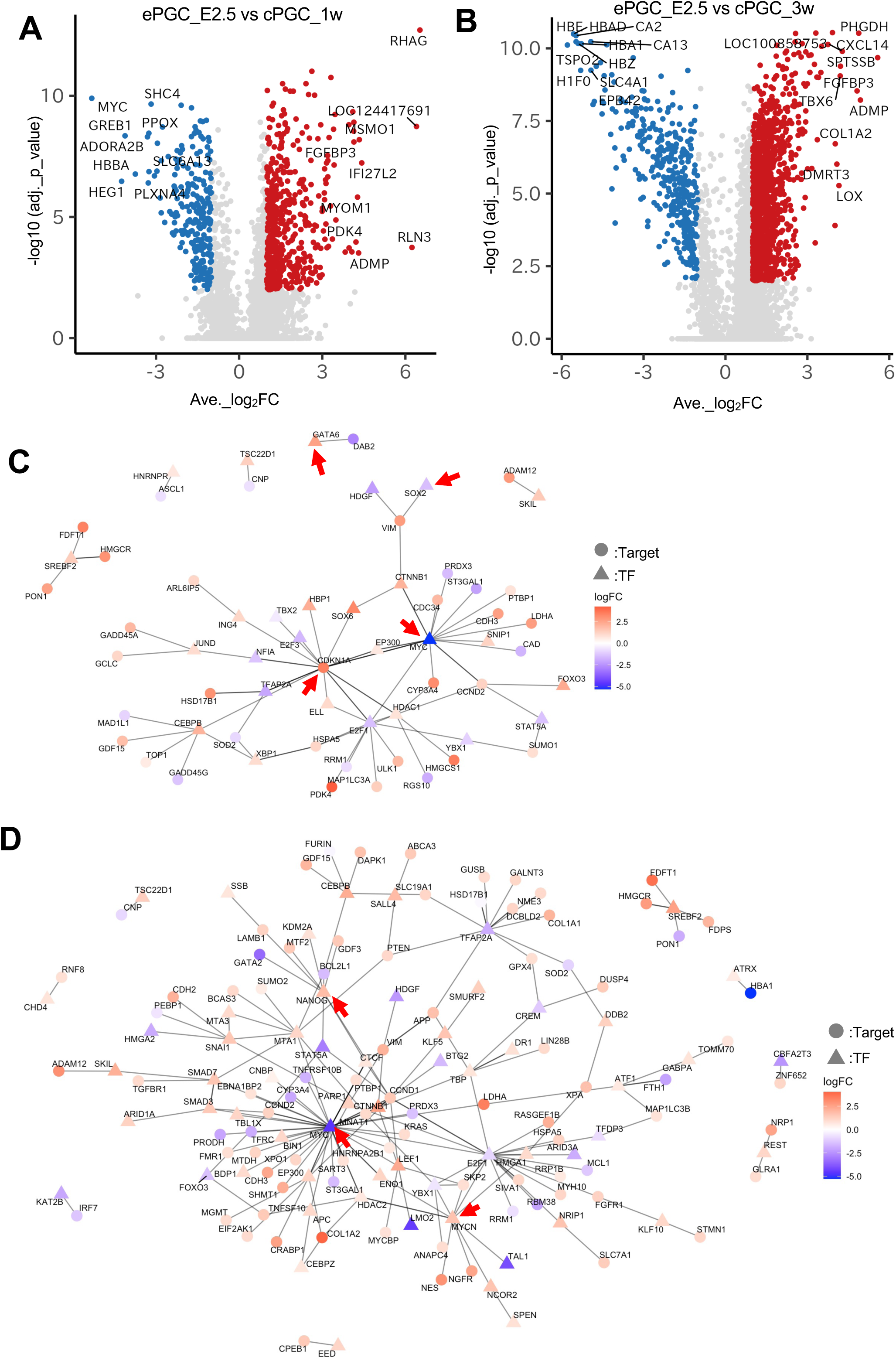
**Gene expression comparison between ePGCs and cPGCs**. (A, B) Volcano plots of differentially expressed genes (DEGs) by comparison between ePGC_E2.5 and cPGC_1w (A), or ePGC_E2.5 and cPGC_3w (B) are shown. In both of those analyses, ePGC_E2.5 samples are set as controls. Red plots indicate the up-regulated genes (average log2FC > 1, adjusted p-value <0.01) and blue plots indicate the down-regulated genes (average log2FC < -1, adjusted p-value <0.01). (C, D) Gene regulatory network (GRN) analysis for DEGs based on the relation between transcriptional factors (TFs, indicated as triangles) and their known target genes (Target, indicated as circles) is shown. (C) GRN analysis for DEGs from comparison of ePGC_E2.5 and cPGC_1w. (D) GRN analysis for DEGs from comparison of ePGC_E2.5 and cPGC_3w. In both C and D, the color value of the nodes indicates the log_2_FC of DEGs. Red arrows in C and D indicate the genes discussed in main texts.

### 4.4 Molecular characterization of PGC-like cells found in embryonic day 0

Finally, we investigated the molecular characteristics of PGC-like cells identified from our LEL-based labeling. Our previous work showed that LEL-labeled PGC-like cells from embryonic day 0 have morphology identical to PGC but are absent from SSEA1(Iikawa *et al*., 2024). Our clustering analysis showed that those PGC-like cells are separately integrated into two clusters, one is absent for DAZL (cluster ePGC_E0) and one is merged into cluster cPGC_3w_mix (Figures 1C, D). We speculated that those results came from the intermediate differentiation state to the germ cell lineage from undifferentiated early embryonic cells. To evaluate this possibility, we check the expression of genes related to cellular pluripotency (OCT4/POU5F3, NANOG, SOX2, KLF4, LIN28A, LIN28B) and germline development (DDX4/VASA, NANOS1, NANOS3, PIWIL1, DAZL, DND1, PRDM14, KIT, TDRD9) (Figure 4, Supp. figure3). We found that the cells in cluster ePGC_E0 showed low level but certain expression of several germline genes (DDX4: Figure 4A, NANOS3: Figure 4B, PIWIL1: Figure 4C, PRDM14: Figure 4E, KIT: Supp. figure 3A, TDRD6: Supp. figure 3B, NANOS1: Supp. figure 3C) with high expression of pluripotent marker genes (Such as OCT4/POU5F3 in Figure 4F). Thus, these data strongly suggest that our identified PGC-like cell population contains intermediate state cells that are supposed to be committed to the germline lineage. Compared to other animal models, the understanding of PGC specification processes in avian species is less well understood since there are no methods to collect PGCs from early embryos. Previous works depend on SSEA1-labeling or fluorescent-tagged DAZL expression for collecting PGCs(Ichikawa & Horiuchi, 2023, Rengaraj *et al*., 2022a, Rengaraj *et al*., 2022b), but both of those markers are absent in our LEL-positive PGC-like cells (Iikawa *et al*., 2024). DDX4/VASA is shown to be the earliest marker for PGC, but a fluorescent-tagged DDX4 transgenic strain is not yet available (Tsunekawa *et al*., 2000). Thus, our results showed that our developed LEL-labeling methods provide a chance to collect PGC-like cell populations those on the PGC specification processes and provide a first chance to understand the molecular mechanism regulating PGC specification in avian species.

**Figure 4.**
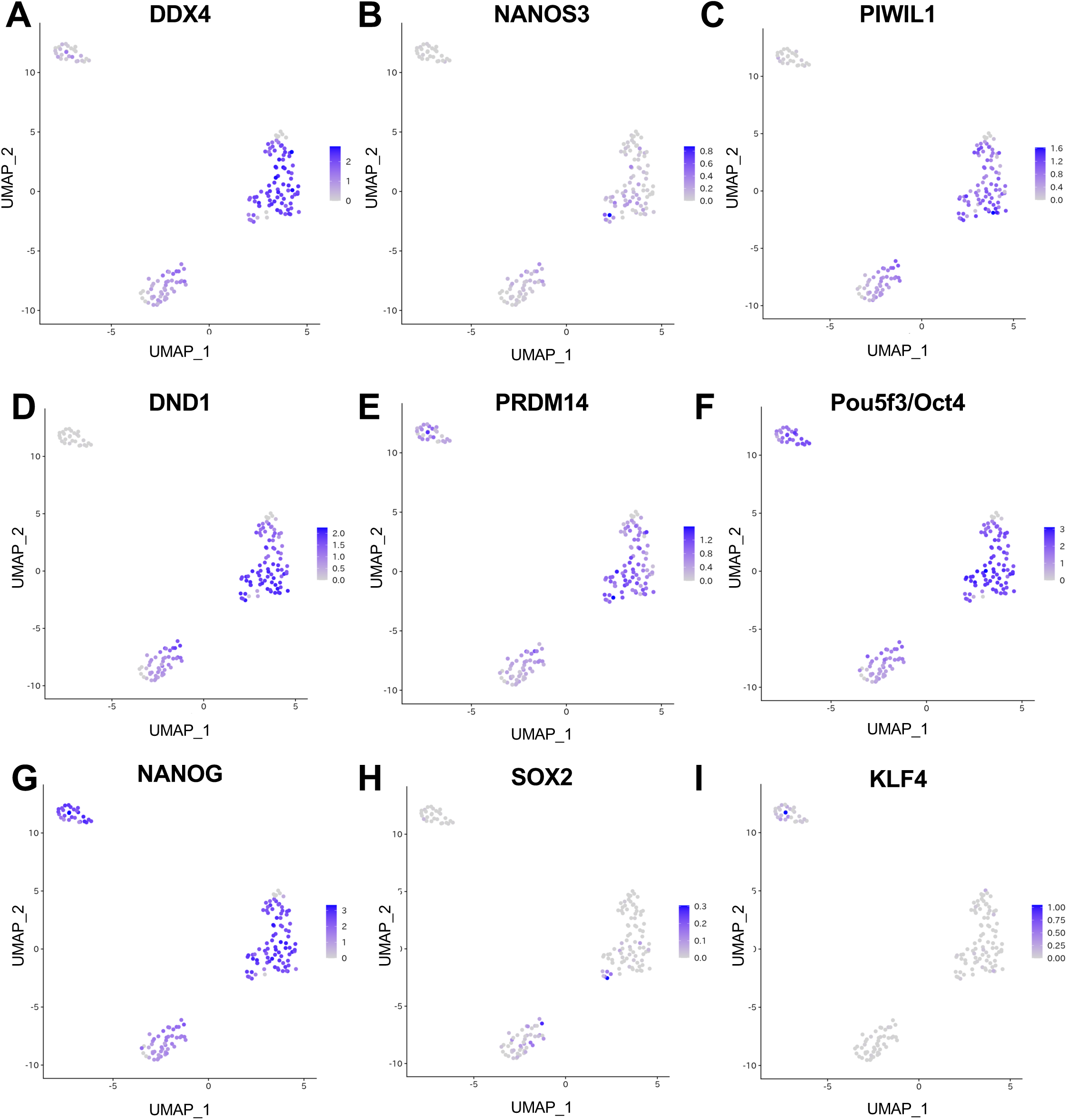
**Molecular characterization of PGC-like cells**. The expression level of indicated germline genes (A-E) and pluripotent marker genes (F-I) in each sample is shown. Color value is based on normalized log-transformed value.

The PGC cultivation system in chickens is valuable for revealing the molecular mechanism regulating unique PGC characters, such as their stem cell-like potential and long-distance migration. In this study, we first provide the molecular evidence to understand the effects of PGC cultivation with SMART-seq-based deep sequencing at the single-cell level. Our study suggests that shifting to MYCN-dependent GRN is essential for the success of PGC cultivation. Also, our study provides the risk of somatic differentiation by cultivation conditions. These results suggest that establishing conditions that simultaneously support reprogramming into a MYCN-dependent gene regulatory network promoting both cell proliferation and maintenance of stem cell identity, while inhibiting somatic differentiation, is essential for successful PGC cultivation. To date, PGC cultivation has not been successful in many animals, including avian species. Thus, our study provides the strategic direction and molecular targets to achieve PGC cultivation in other avian species. During the preparation of this manuscript, Doddamani and colleagues reported the long-term cultivation method of goose PGCs (Doddamani *et al*., 2025). The study revealed that Activin A and FGF2 have opposite effects on goose PGC in vitro propagation; Excessive Activin A inhibits propagation of PGCs (Doddamani *et al*., 2025). This observation is consistent with our results suggesting that Activin A signaling through SMAD3 function suppresses MYC-dependent GRN, and emphasizes the importance of balancing the activity of three major constituents, Activin A, FGF2 and Insulin. In addition to the molecular differences between ePGCs and cPGCs, our study provides molecular evidence suggesting that our LEL-labeling method is a useful tool for collecting PGCs from the early stage of the chick embryo. Although the specification process of PGC is a fundamental question of developmental biology and is known to be diverse among animal species, PGC specification processes in avian species are less understood because of the lack of molecular information. Our study suggests that the LEL-positive PGC-like cells at embryonic day 0 are a partially differentiated cell population in the germline. Thus, our results and developed methods could be a key to understanding the PGC specification processes in avian species. Therefore, our SMART-seq-based scRNA-seq analysis provides fundamental knowledge not only for the future success of PGC cultivation in non-chicken animals but also for understanding the PGC specification processes in avian species.

## 3. Materials & Methods

### 3.1 Animals

Fertilized eggs from the BL-E strain of brawn leghorn chicken were provided by Nagoya University through the National Bio-Resource Project of the MEXT, Japan. The eggs were incubated at 38.5lJ in a high-humidity chamber (50-60%). Chicken embryos at stages E0 and E2.5 were dissected and collected after 30 minutes and 60 hours of incubation, respectively. All animal experiments were conducted in accordance with the ethical guidelines for animal experimentation established by Kyushu University.

### 3.2 PGC culture

PGCs collected from E2.5 chicken embryos were cultivated as described previously(Iikawa *et al*., 2024). In brief, the PGCs were cultivated in modified FAcs medium at 38.0lJ with 5% CO_2_ in humidity incubator until the investigated time points (1 week or 3 weeks). The FAcs medium composition includes; 65.5% DMEM (high glucose, Ca^2+^-free, no-glutamine) (Nakarai tesque, Japan), 21.8% DDW (Fuji film, Japan), 1x Nucleotides (Merck-Millipore, U.S.A.), 2.0mM GlutaMax [Thermo Fisher Scientitic (Thermo), U.S.A.), 1.2mM Sodium Pyruvate (Thermo), 1x NEAA (Thermo), 55µM beta-mercaptoethanol (Fuji film), 1% Penicillin-Streptomycin-Amphotericin (PSA) (Fuji film), 2% B27 Supplements (Thermo), 0.2% Chicken Serum (Biowest, France), 0.1mg/ ml Heparin (Sigma-Aldrich, U.S.A.), 2mg/ ml Ovoalbmin (Sigma-Aldrich), 25ng/ ml Activin A (API, Japan), 4ng/ml FGF Chimera (Fuji film), 100 µM CaCl_2_ (Nakarai tesque)(Whyte *et al*., 2015). Half of the medium was replaced with fresh medium every 2 days.

### 3.3 PGC collection

PGCs from E0 and E2.5 embryos were collected by labeling with *Lycopersicon Esculentum* (Tomato) lectin conjugated with DyLight^®^488 (DL-1174, Vector Laboratories, U.S.A.) by Fluorescence-Activated Cell Sorting (FACS) (SH800, Sony, Japan) as previously described (Iikawa *et al*., 2024). Sorted cells were collected in a 96-well plate (Ro-bind, 0030 129.504, Eppendorf, Germany) filled with Lysis Buffer containing RNase inhibitor (Components of kit described below, TAKARA, Japan) and immediately frozen with dry ice. The lysed cells were used to prepare the libraries as described below.

### 3.4 SMART-Seq

The SMART-Seq libraries were prepared with SMART-Seq Single Cell PLUS Kit (TAKARA) following the manufacture’s instruction. The libraries were sequenced by NovaSeq 6000 with 150 bp pair-end sequencing (Illumina).

### 3.5 Data analysis

The treated reads were mapped with chicken genome (bGalGal1.mat.broiler.GRCg7b). The cluster analysis was performed with Seurat software (5.1.0) on R (4.4.1). GO term enrichment analysis was performed with clusterProfiler (4.15.1) on R(4.5.0), and enrichment analysis to identify tissue-specific gene expression signatures in the DEGs was performed with enrichR (3.4) with ARCHS4_Tissues gene sets on R(4.5.0). Network analysis was performed with igraph (2.1.4) on R(4.5.0). The network data was obtained from TRRUST version2 (human), and chick differentially expressed genes were converted to human ortholog by using g:Profiler web tool.

### 3.6 Data availability

The sequencing data have been deposited in the DDBJ under accession number PRJDB17925.

## 4. Acknowledgement

This work was supported in part by Grants-in-Aid for Scientific Research from Japan Society for Promotion of Science (JSPS) (KAKENHI Grant Number: 23K05777, 23K17933 to Y.H., and 22H02634, JP22H04925 (PAGS) to D.S.

## 5. Author contributions

D.S. designed the experiments. H.I., Y.S. and A.K. performed the experiments. Y.H., A.D. and H.H. analyzed the data. Y.H. and D.S. wrote the manuscript. All authors discussed the results and commented on the manuscript.

---

**Supp. Figure 1.**
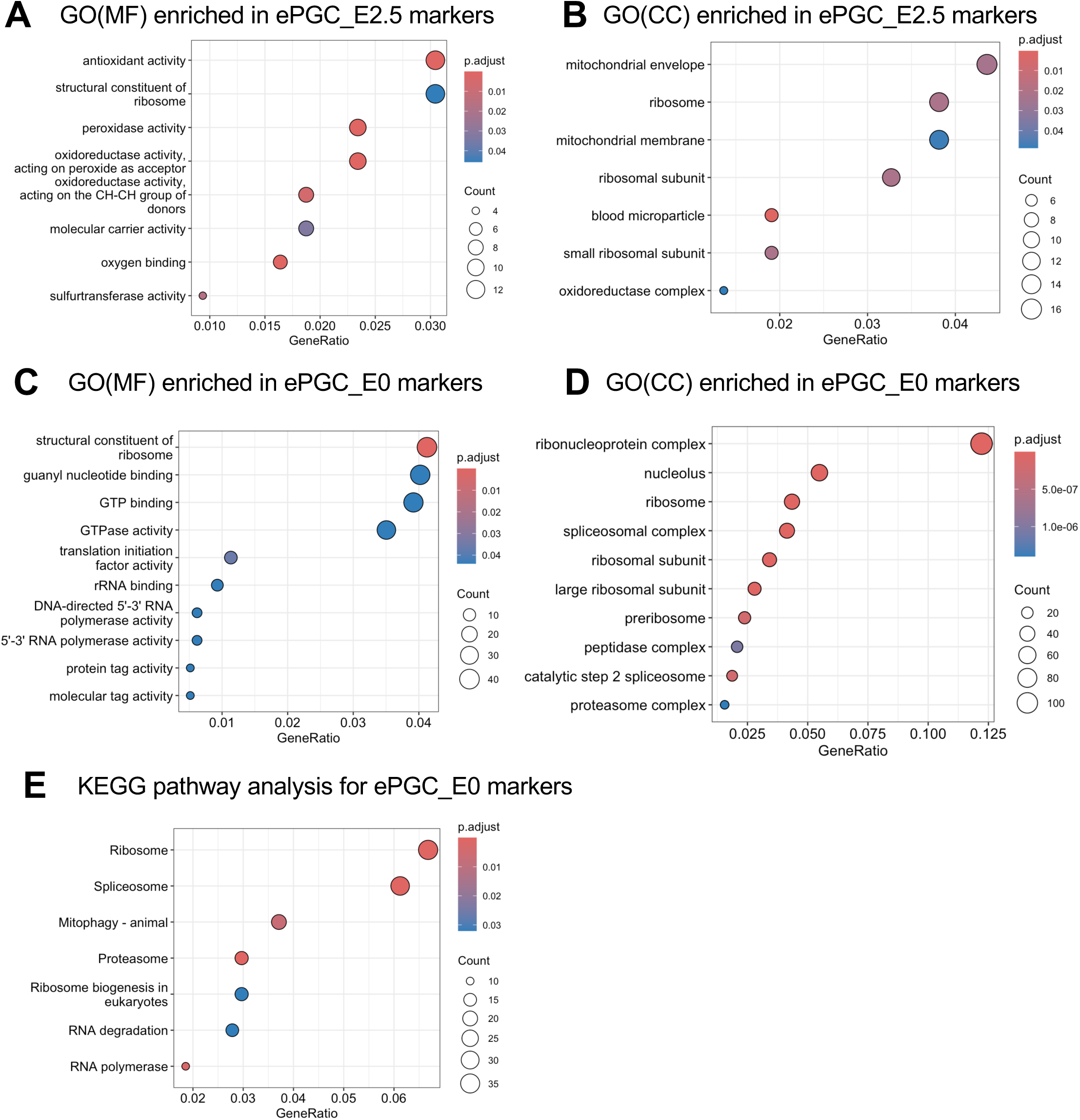
Molecular feature of PGC clusters. The results of Gene Ontology (GO) enrichment analysis or KEGG pathway analysis for marker genes of each cluster are shown. The analyzed marker gene source and type of analysis are indicated at the top of each figure (GO(MF); GO enrichment analysis for Molecular Function (MF) category, GO(CC); GO enrichment analysis for Cellular Components (CC) category).

**Supp. Figure 2.**
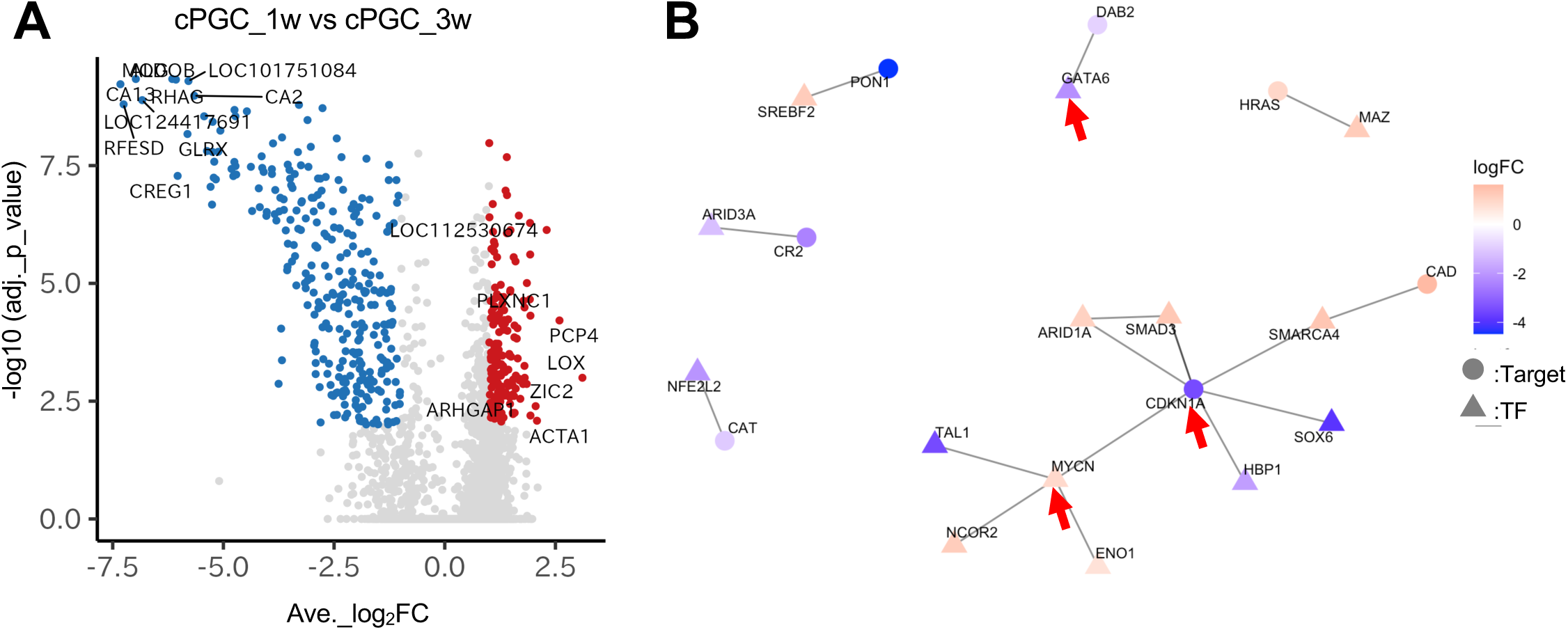
**Gene expression comparison between cPGC_1w and cPGC_3w**. (A) Volcano plots of differentially expressed genes (DEGs) by comparison between cPGC_1w and cPGC_3w are shown. cPGC_1w sample is set as controls. Red plots indicate the up-regulated genes (average log2FC > 1, adjusted p-value <0.01) and blue plots indicate the down-regulated genes (average log2FC < -1, adjusted p-value <0.01). (B) Gene regulatory network (GRN) analysis for DEGs between cPGC_1w and cPGC_3w based on the relation between transcriptional factors (TFs, indicated as triangles) and their known target genes (Target, indicated as circles) is shown. Red arrows in B indicate the genes discussed in main texts.

**Supp. Figure 3.**
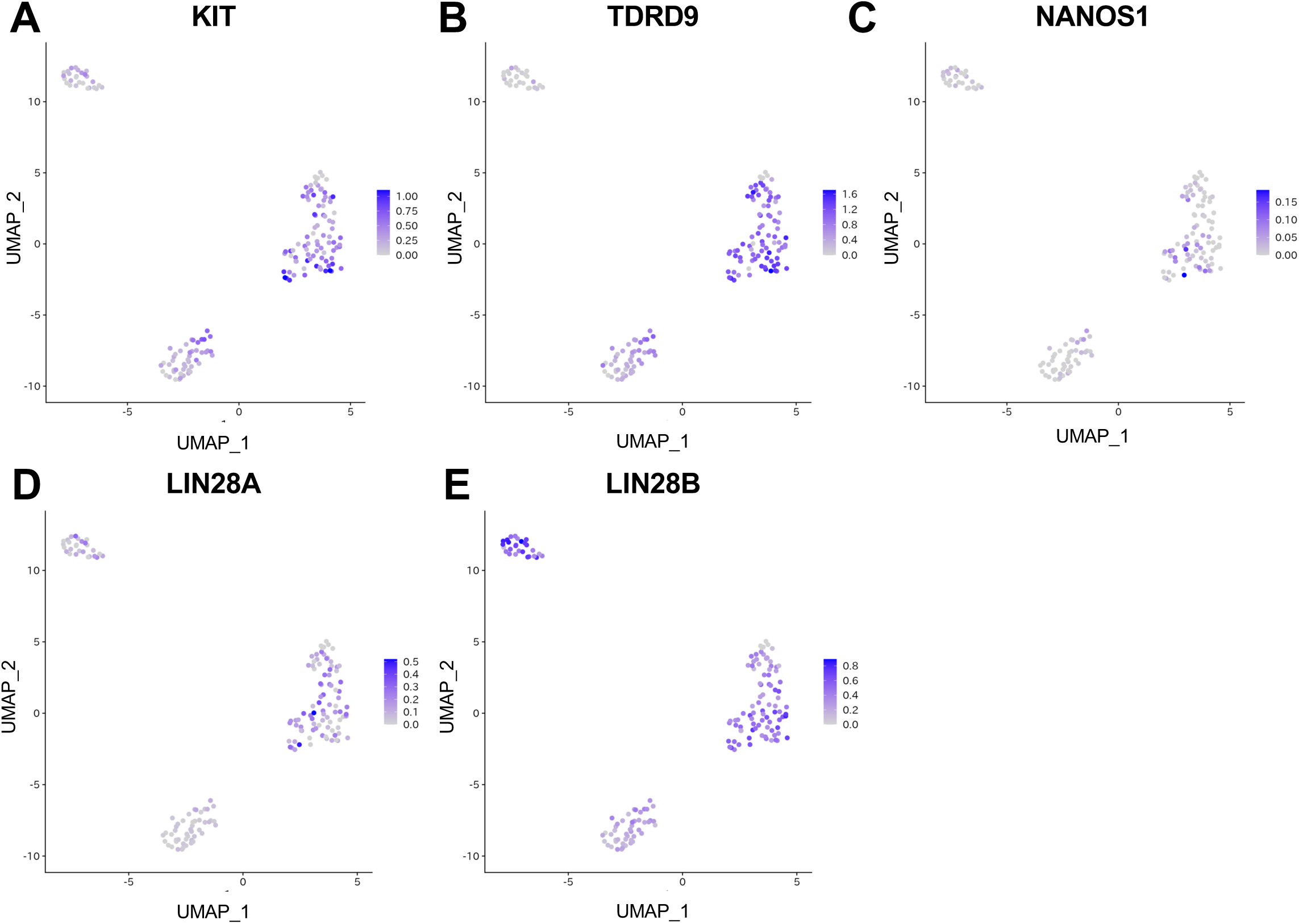
**Molecular characterization of PGC-like cells**. The expression levels of indicated germline genes (A-C) and pluripotent marker genes (D, E) in each sample are shown. Color value is based on normalized log-transformed value.

